# Mesolimbic local field potentials are modulated by motor control

**DOI:** 10.1101/2024.07.02.601451

**Authors:** Leah G Mann, Helen Qian, Natasha Cindy Hughes, Zixiang Zhao, Balbir Singh, Zhengyang Wang, Graham W. Johnson, Jenna N. Fulton, Bailu Yan, Hakmook Kang, Rui Li, Benoit M. Dawant, Dario J. Englot, Christos Constantinidis, Shawniqua Williams Roberson, Daniel O. Claassen, Sarah K. Bick

## Abstract

**Background and Objectives:** While historically cortico-basal ganglia-thalamocortical loops were believed to process limbic and sensorimotor data in parallel, there is now evidence to suggest that the two information streams can be processed in a single open loop. However, the limbic-motor interface remains insufficiently characterized. We sought to further investigate how extrastriatal regions may regulate motor output by examining electrophysiological activity in these areas during a response inhibition paradigm.

**Methods:** We recorded local field potentials (LFPs) from epilepsy patients implanted with intracranial depth electrodes for seizure localization purposes. Participants performed the stop-signal task, during which they made speeded choice reactions to “go” stimuli and occasionally inhibited their reactions in the incident of a “stop” signal. To compare power during movement and the absence of movement, we applied a Wilcoxon signed-rank test. Additionally, we performed a linear mixed-effects model to relate power in limbic regions to power in the motor cortex. Finally, we implemented exploratory analyses to identify power differences for correct go versus correct stop trials and for correct stop versus incorrect stop trials using cluster-based permutation testing.

**Results:** 14 patients participated. A comparison between movement and baseline fixation revealed that motor response is associated with reduced beta (15-35 Hz) power in the amygdala, hippocampus, and motor cortex and reduced gamma (35-100 Hz) power in the amygdala and hippocampus. Moreover, average beta and gamma power in the amygdala and hippocampus during motor execution were positively associated with average beta and gamma power in the motor cortex. Additionally, we identified significant differences between correct go and stop trials in delta (1-4 Hz) power for all three regions and in theta (4-8 Hz) power for the amygdala and motor cortex. Likewise, we identified significant differences between correct and incorrect stop trials in delta power for the hippocampus and motor cortex, in theta power for the motor cortex, in alpha (8-15 Hz) and beta power for the amygdala and motor cortex, and in gamma power for all three areas.

**Discussion:** These correlations between neural oscillations in the hippocampus and amygdala and movement strengthen the notion of mesolimbic modulation of motor activity.

## Introduction

Connections between the limbic and motor pathways have long been suspected despite the unique information carried and functions performed by these different networks. The ventral striatum is frequently recognized as the interface between these two discrete systems, receiving limbic efferents and converting the signals into motor output.^1,2^ Evidence for this coupling arises from movement disorders, such as Parkinson’s disease, that often co-exist with impaired inhibitory control. For these patients, differences in action control and movement have been correlated with differences in dopamine receptor availability in mesial temporal regions, which are primary components of the limbic system.^3,4^ Moreover, functional movement disorders provide even more pertinent evidence of a limbic-motor connection. These syndromes are characterized by unusual movements that are likely a consequence of psychological underpinnings and have been associated with enhanced amygdala responsiveness and greater functional connectivity between the amygdala and supplementary motor area.^5–8^ Beyond clinical evidence, animal histology and electrophysiology studies have demonstrated the influence of limbic information on the motor loop through a pathway from ventral striatum to substantia nigra to motor thalamus.^9,10^

Despite the progress made, research on the activity and mechanisms of the functional limbic-motor loop in humans has been stagnated by techniques with poor temporal or spatial resolution. Functional imaging techniques measure blood flow rather than direct neural activity and as such have limited temporal resolution. Scalp electroencephalography (EEG) has excellent temporal resolution but poor spatial resolution, particularly for deep structures. Stereoelectroencephalography (sEEG) recordings performed in medically refractory human epilepsy patients for the purpose of seizure localization allow direct local field potential recordings from deep structures, including the hippocampus and amygdala, with excellent spatial and temporal resolution. As such, sEEG provides an optimal opportunity to study limbic and motor systems in tandem.

It is widely accepted that sensorimotor beta (15-35 Hz) power decreases during motor planning and execution and rebounds following movement cessation.^11–13^ Studies have also demonstrated analogous reductions in motor cortex alpha (8-15 Hz) power with movement production.^14–16^ On the other hand, gamma (35-100 Hz) power in the motor cortex has been shown to increase during motor preparation.^17–20^ Yet, whether similar changes in beta, alpha, or gamma oscillations in the amygdala and hippocampus are also present relative to movement has not been resolved. Additionally, it is probable that mesolimbic power in other frequency bands plays a role in movement-related behaviors.

In line with an integrated limbic-motor loop, we hypothesized that neural oscillations in the amygdala and hippocampus contribute to movement and that limbic beta oscillations would be associated with motor output. To test this, we examined local field potential (LFP) recordings from the amygdala, hippocampus, and motor cortex as epilepsy patients with implanted sEEG electrodes performed a stop-signal task. The action control paradigm is ideal for investigating power changes accompanying an absence of movement, a presence of movement, and an inhibition of movement. We describe amygdala and hippocampus movement-coupled power changes in beta and gamma bands and additional changes in delta power when examining inhibitory instances.

## Methods

### Participants

Epilepsy patients (n = 14, sex = 9F/5M, age range = 29-59 years) undergoing sEEG were recruited from Vanderbilt University Medical Center. Patients had previously been diagnosed with medically intractable epilepsy and were undergoing sEEG electrode implantation for clinical seizure localization. All participants were screened to confirm that they met inclusion and exclusion criteria. Participants were omitted from entering the study if they were under 18 years of age or were unable to understand the task. Table 1 presents demographic and clinical information. Electrodes were implanted under general anesthesia, with the number and locations of electrodes individualized for each patient according to clinical considerations. After electrode implantation surgery, patients were admitted to the epilepsy monitoring unit where they remained for continuous seizure monitoring. Here, antiepileptic medications were gradually tapered. While participants were monitored, we collected local field potentials from the sEEG electrodes using the Natus data acquisition system (Natus, Middleton, WI).

**Table 1.** Demographic and clinical evaluation of the participants.

| **Participant ID** | **Sex** | **Handedness** | **Seizure Onset** |
| --- | --- | --- | --- |
| 1 | F | Left | Left occipital and left posterior temporal nodular heterotopias |
| 2 | M | Left | Right mesial temporal |
| 3 | F | Right | Left mesial temporal |
| 4 | F | Right | Right mesial and basal temporal region |
| 5 | M | Right | Left mesial temporal |
| 6 | F | Right | Left anterior cingulate and orbitofrontal |
| 7 | M | Right | Left and right mesial temporal, right anterior cingulate |
| 8 | F | Right | Left mesial temporal |
| 9 | F | Right | Bilateral mesial temporal |
| 10 | F | Left | Bilateral mesial temporal, right periventricular nodular heterotopia |
| 11 | F | Right | Right mesial temporal |
| 12 | M | Right | Right mesial temporal |
| 13 | F | Right | Left temporal cortex |
| 14 | M | Right | Left mesial temporal, right frontal/insula |

### Standard Protocol Approvals, Registrations, and Patient Consents

The study was carried out in accordance with the Declaration of Helsinki, and all subjects provided written, informed consent before participating in the study in compliance with the standards of ethical conduct in human investigation regulated by the local Institutional Review Board.

### Stop-signal task

Patients performed a manual version of the stop-signal task using a portable, 13-inch tablet (Microsoft Surface, Microsoft Corporation, Redmond, WA).^21^ Participants carried out the task in their rooms in the epilepsy monitoring unit as neural recordings were continuously captured. The task was executed through MATLAB 2022 (MathWorks, Natick, MA) and the Psychophysics Toolbox Version 3 core.^22^ Before proceeding, patients were provided with task directions and instructed to make their responses as quickly and accurately as possible.

The stop-signal task consists of two trial types, namely Go trials and Stop trials, that are presented at random to participants. Each trial began with a fixation point that remained on the screen for 1500 ms. Go trials constituted the majority of trials and involved the appearance of a left- or right-pointing gray arrow as the “go” signal. Patients were asked to press a button in accordance with the direction of the go arrow, making a right-hand button press when they were shown a right-pointing arrow and a left-hand button press when they were shown a left-pointing arrow. The speed with which a participant made a response after the appearance of the go stimulus was measured as go reaction time (GoRT). The arrow disappeared following a button press or once 1200 ms had elapsed (Figure 1).

**Figure 1.**
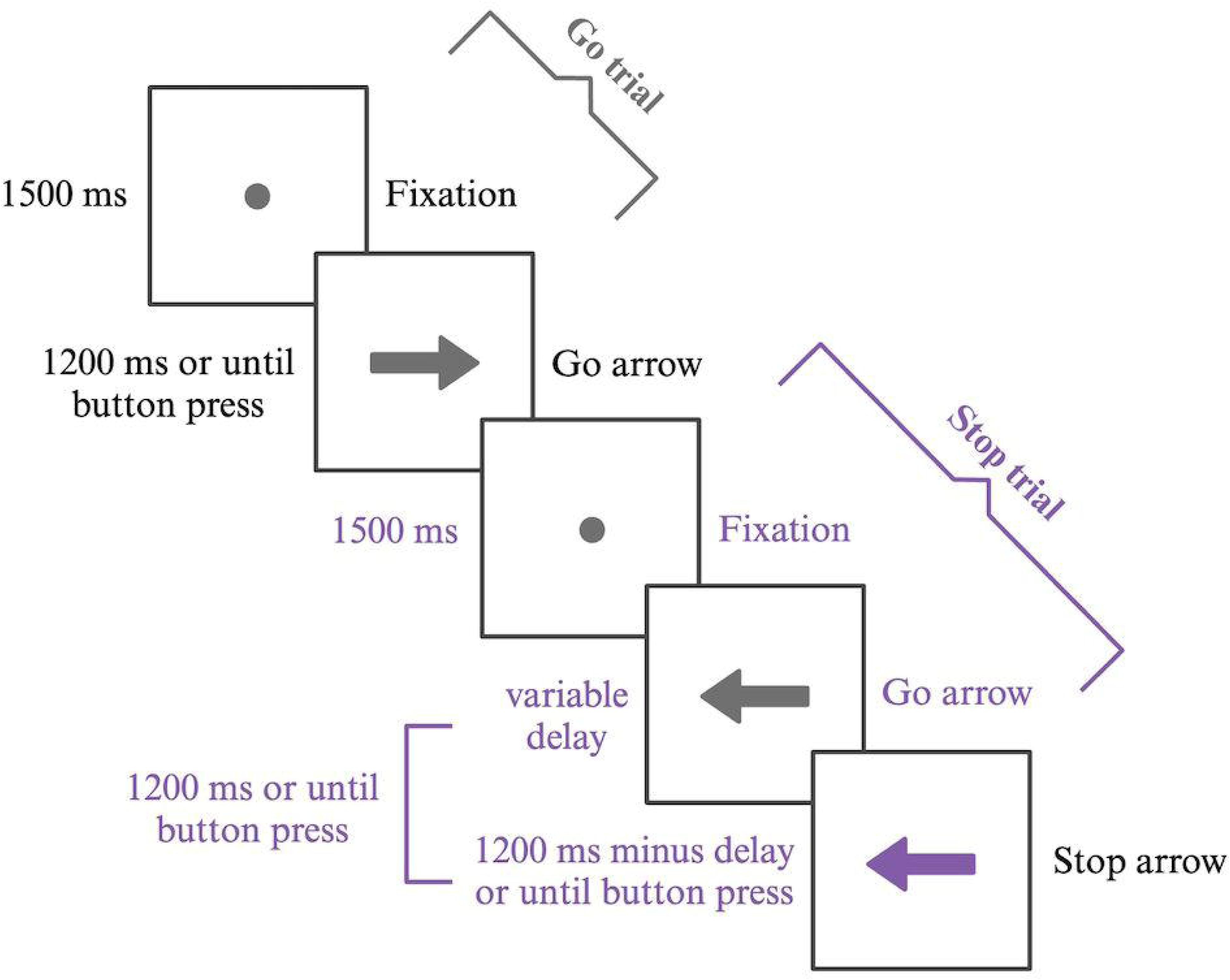
Stop-signal task design. Example of the two trial types in the stop-signal task. 75% of the trials were go trials, during which a gray arrow appeared and participants were to make a button press according to the direction of the arrow. 25% of the trials were stop trials, during which a gray arrow appeared but was soon replaced by a purple arrow. On these trials, participants were to inhibit a button press.

About 25% of the total trials were stop trials and involved the go arrow changing color to purple soon after appearing on the screen. Patients were to inhibit their response and avoid pressing a button once they perceived the purple arrow as the “stop” signal. The time between the occurrence of the go arrow and its color change differed from trial to trial according to a staircase tracking method based on performance.^23^ The delay for the first stop trial of each block was set at 200 ms and was subsequently adjusted after each stop trial, increasing by 50 ms if the patient successfully inhibited a response and decreasing by 50 ms if the patient erroneously made a response. Each administration of the stop-signal task involved 10 practice trials, followed by two blocks of 75 experimental trials. The design of the stop-signal task is shown in Figure 1.

When a go signal, stop signal, or response was initiated, a transistor-transistor logic (TTL) pulse was sent to the trigger channel of the EEG amplifier in order to allow alignment of task behavioral events with sEEG recordings.^24^ To analyze task performance, the mean method was applied for estimating mean stop-signal reaction time (SSRT).^21,25^

### Electrode localization

The intracranial electrodes implanted for neural recordings measured 0.8 mm in diameter and contained 8-16 contacts that were spaced between 2.5– 4.3 mm center-to-center (PMT Corporation, Chanhassen, MN). Each patient underwent an MRI scan prior to sEEG implantation surgery and a CT scan following surgery with electrodes in place. The images from these preoperative and postoperative scans were coregistered to identify coordinates of electrode contacts using the CRAnial Vault Explorer (CRAVE) software.^26^ The FreeSurfer software package (http://surfer.nmr.mgh.harvard.edu) was then used to map electrode coordinates to specific anatomic brain regions according to the Desikan-Killiani Atlas. Mapping between electrode channels and brain regions was further confirmed by an epileptologist (SWR).

### Neurophysiology processing

LFP recordings were processed and analyzed using MATLAB (MathWorks, Natick MA) and the FieldTrip toolbox software.^27^ Raw LFP signals from contacts within the brain regions of interest (amygdala, hippocampus, and motor cortex) were visually inspected for artifacts. Channels and trials that contained excessive artifact were excluded from further analysis. Additionally, electrode channels were excluded from analysis if located within a patient’s seizure onset zone, defined as an area with first ictal changes as determined by an epileptologist (SWR). As each electrode contained multiple contacts and electrodes were frequently positioned in bilateral regions, many subjects had multiple electrode channels within each region of interest.

LFP recordings were performed with a sampling rate of 512 Hz. Data was high-pass filtered at 1 Hz and low-pass filtered at 200 Hz. Additionally, a band-stop filter was applied to remove 60 Hz line noise. TTL pulses were used to align task execution information with LFP recordings. Data was divided into trials and aligned to the appearance of the fixation point, the appearance of the go signal, the appearance of the stop signal, or the response. Spectral power was calculated using a multitaper time-frequency transformation and a Hanning taper. Spectral power was then z-scored across all trials for each channel and frequency to allow comparison across subjects and channels.

### Statistical analyses

To examine movement-related limbic and motor cortex oscillation changes, we assessed power during the execution of a motor response in relation to power during the absence of a motor response. To evaluate our hypothesis that limbic and motor beta power would be modulated by movement, we averaged beta power within each channel during a 500 ms response period from 250 ms before a response was made to 250 ms after a response was made for correctly performed go trials. Similarly, we averaged beta power within each channel during a 500 ms baseline fixation period from 750 ms to 1250 ms following the appearance of the fixation point. We then applied the Wilcoxon signed-rank test to compare beta power during these two task epochs for all three regions. The Bonferroni correction, with alpha level set to 0.05, was used to correct for multiple comparisons. We completed the same procedure to compare gamma oscillations during a correct go response to gamma oscillations during fixation.

Furthermore, we analyzed the relationship between motor cortex and limbic region power during responding on a single-trial level. To do this we averaged beta power across all motor cortex channels for a 500 ms response period from 250 ms before response to 250 ms after response for each correct go trial for each participant. Only participants with electrodes in both the motor cortex and limbic regions were included in this analysis. We then averaged beta power across all amygdala channels and hippocampus channels for the same time interval for each trial for each patient. Next, we performed two linear mixed-effects models with subject-specific random intercepts and imposed first-order autoregressive, AR(1), correlation structures to compare beta power in the motor cortex with beta power in the amygdala and hippocampus. For these models, beta power in the amygdala or hippocampus served as the dependent variable, beta power in the motor cortex as the independent variable, and age and sex as covariates. We completed the same procedure to compare gamma power in the motor cortex with gamma power in the amygdala and hippocampus.

To probe the possibility that limbic and motor region power fluctuations are associated with motor inhibition we performed exploratory analysis to identify power differences between correct go and correct stop trials. We first identified power differences between correct go and stop trials in the motor cortex and then evaluated whether power in the same frequency bands and time ranges was also significantly different between correct go and correct stop trials in limbic regions. To do this, we first averaged power aligned to the go arrow appearance within the delta (1-4 Hz), theta (4-8 Hz), alpha (8-15 Hz), beta (15-35 Hz), and gamma (35-100 Hz) bands in motor cortex channels and averaged all correct go and correct stop trials for each channel. Next, we performed cluster-based permutation testing to identify time clusters with significantly different power between correct go and correct stop trials in the motor cortex for a 600 ms time window beginning 150 ms after the go arrow onset.^28^ This time window was selected to account for both average GoRT and average SSRT. For each frequency band that was associated with a significant cluster in the motor cortex, we averaged power during the significant time period within each channel in the amygdala and hippocampus for correct go and correct stop trials separately. We then applied the Wilcoxon signed-rank test to compare power during this time range between these two trial types for the limbic regions. Finally, we applied the Bonferroni correction to correct for multiple comparisons.

We conducted a second exploratory analysis to investigate whether differences in oscillatory power between correct stop and incorrect stop trials aligned to the appearance of the stop signal are seen in limbic regions. We again first identified power changes between these two conditions in motor cortex and then evaluated whether these were also present in limbic regions. We used cluster-based permutation tests for each power band to identify time clusters that were significantly different between stop trials in which movement was correctly inhibited and those in which it was incorrectly executed in the motor cortex. We analyzed power in the motor cortex during a 2000 ms time window following the appearance of the stop arrow. Again, we limited our analyses for the amygdala and hippocampus to the time intervals determined to have significant power differences between correct and incorrect stop trials in the motor cortex. We averaged power in the relevant band for each time interval and applied the Wilcoxon signed-rank test followed by the Bonferroni correction for multiple comparisons.

### Data Availability

Data reported in this article will be made available by the lead contact upon request from a qualified investigator.

## Results

All 14 participants completed the stop-signal task while neurophysiology recordings were collected. Twelve patients had recordings from the amygdala (71 total channels), with 5 of them having electrodes in bilateral structures. Nine patients had recordings from the hippocampus (68 total channels), with 4 of them having electrodes in bilateral structures. Six patients had recordings from the motor cortex (35 total channels), with 1 of them having electrodes in bilateral structures. Participant demographic and clinical information is shown in Table 1.

The average GoRT was estimated as 625.91 ± 128.92 (± SD) ms and the average SSRT was estimated as 285.84 ± 61.28 ms. Supplementary Table 1 presents each patient’s behavioral parameters for task performance as calculated by the mean method.

### Beta and gamma power changes in limbic and motor regions during movement

Given the known role of beta and gamma power in modulating movement, we first evaluated whether oscillatory power in these frequency ranges in the amygdala, hippocampus, and motor cortex was modulated by movement. Figure 2A shows the spectrograms for correct go trials aligned to response, illustrating average z-scored power in all frequency bands. Figure 2B-C depict the time courses of changes in beta and gamma power aligned to response during correct go trials.

**Figure 2.**
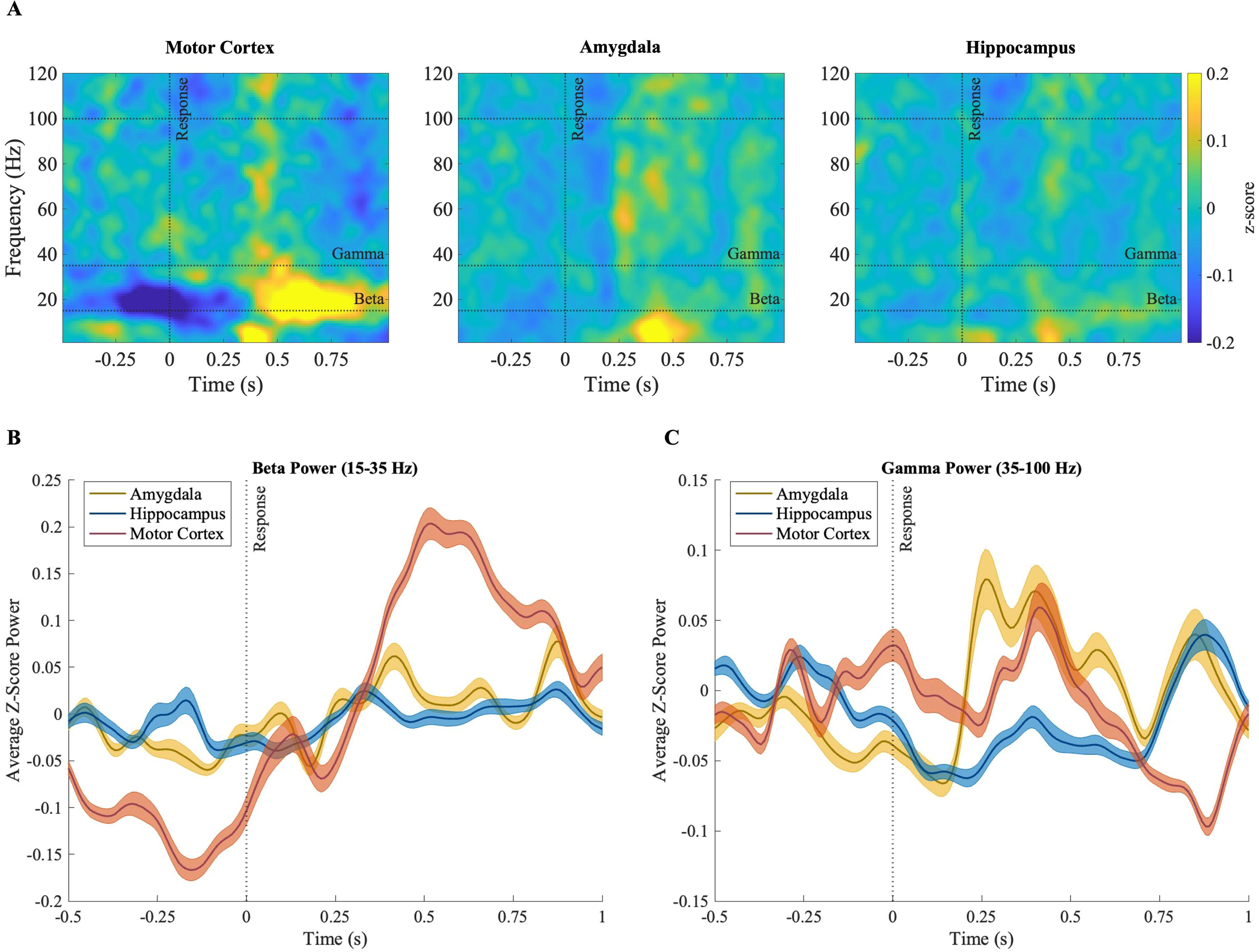
Time course of power during correct go trials. Average z-scored power for correct go trials aligned to response during the stop-signal task in the (A) motor cortex, amygdala, and hippocampus. (B) Average power in the beta frequency range (15-35 Hz) for correct go trials aligned to response in the motor cortex, amygdala, and hippocampus. (C) Average power in the gamma frequency range (35-100 Hz) for correct go trials aligned to response in the motor cortex, amygdala, and hippocampus. Shading around lines indicates 99% confidence intervals.

Comparing beta power during movement with beta power during fixation, we found a significant difference for all regions (Wilcoxon-signed rank test, motor cortex: *p* = 7.9×10^−5^, *z* = −4.36; amygdala: *p* = 1.1×10^−7^, *z* = −5.62; hippocampus: *p* = 2.5×10^−4^, *z* = −4.1), with motor response associated with reduced average beta power compared to the baseline fixation period. Similarly, comparing gamma power during these two task epochs demonstrated a significant difference for the amygdala and hippocampus (Wilcoxon-signed rank test, amygdala: *p* = 1.7×10^−5^, *z* = −4.68; hippocampus: *p* = 3.6×10^−9^, *z* = −6.19), with motor response associated with lower average gamma power compared to baseline. The difference in motor cortex gamma between movement and fixation was not significant (Figure 3).

**Figure 3.**
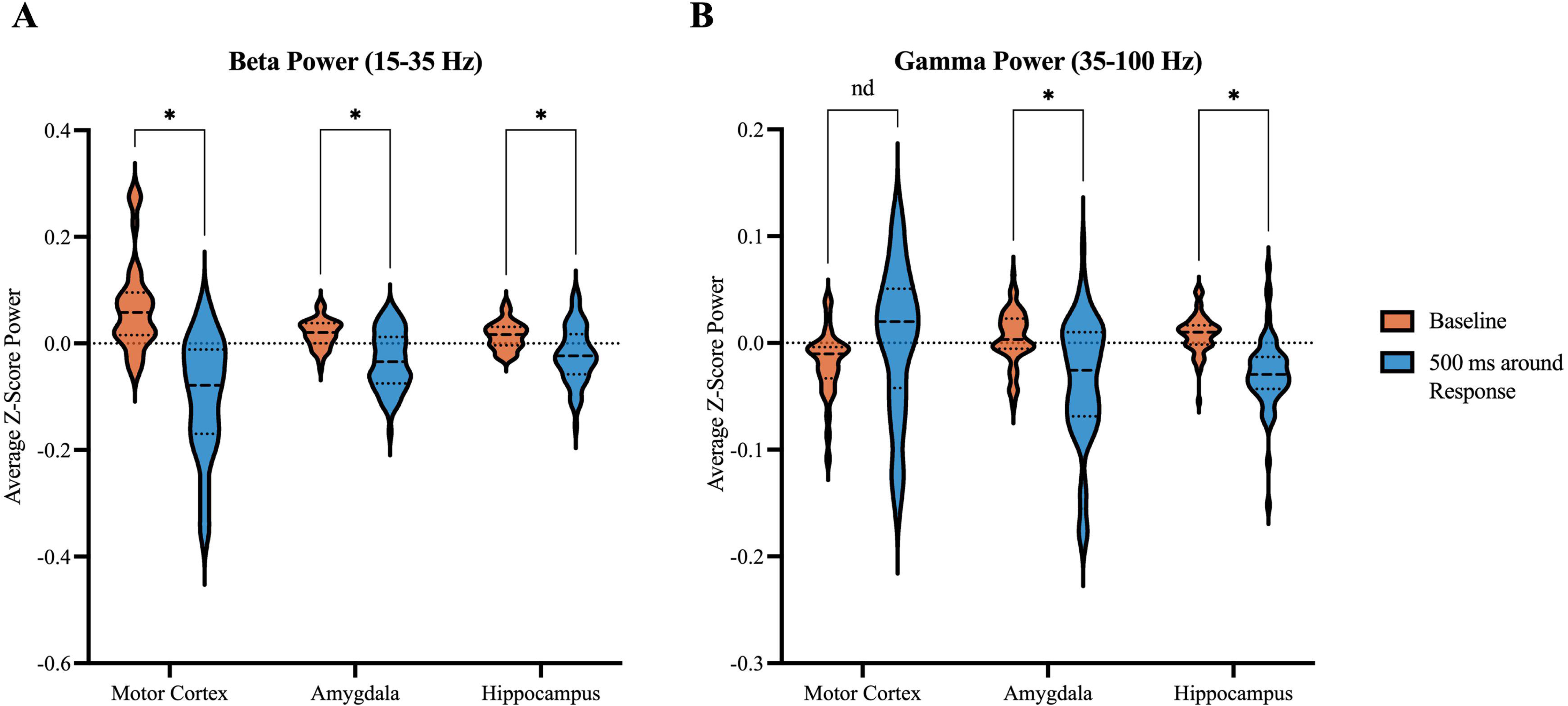
Beta and gamma power during baseline compared to 500 ms around motor response. Violin plots comparing z-scored power during fixation (time period of 750 ms to 1250 ms following appearance of fixation point) and during a correct go response (time period of 250 ms before response to 250 ms after response) in the motor cortex, amygdala, and hippocampus. (A) Violin plot of z-scored power in the beta frequency range (15-35 Hz) during these two task epochs (*p* < 0.01 for all regions). (B) Violin plot of z-scored power in the gamma frequency range (35-100 Hz) during these two task epochs (*p* < 0.01 for amygdala and hippocampus).

### Correlations between power in limbic and motor regions during motor response

The linear mixed-effects model revealed a significant positive relationship between beta power in the motor cortex and beta power in the amygdala and hippocampus on a trial by trial basis (linear mixed-effects model with imposed correlation structure, amygdala: *p* < 0.001, coefficient = 0.25; hippocampus: *p* < 0.001, coefficient = 0.39) during the execution of a motor response. Similarly, we found a significant positive relationship between gamma power in the motor cortex and gamma power in the amygdala and hippocampus (linear mixed-effects model with imposed correlation structure, amygdala: *p* < 0.001, coefficient = 0.50; hippocampus: *p* < 0.001; coefficient = 0.45) during the execution of a motor response (Figure 4).

**Figure 4.**
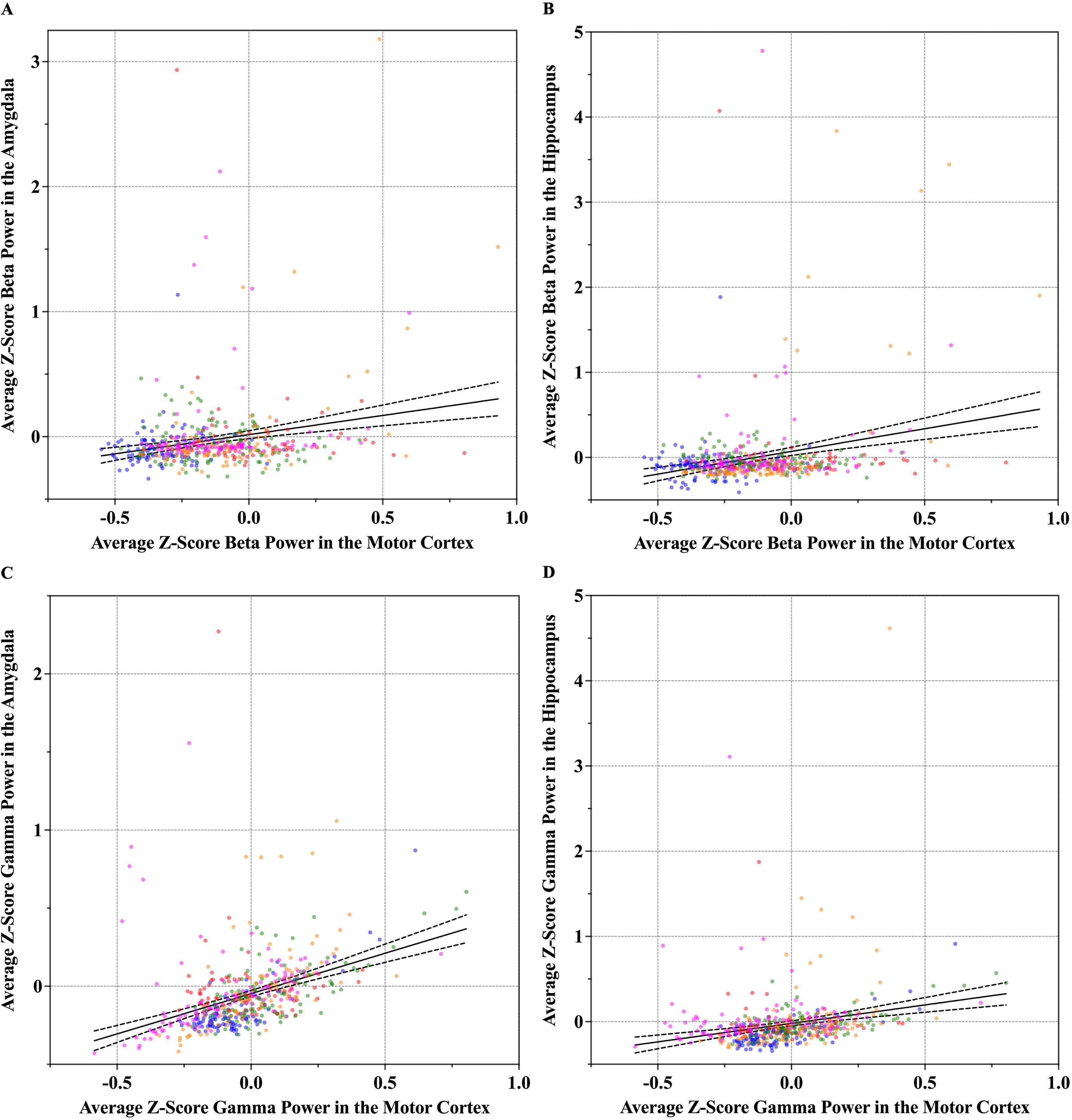
Relationship between power in limbic regions and power in the motor cortex. Scatterplots with lines of best fit displaying the relationship between power in the amygdala and hippocampus and power in the motor cortex during correct go responses (time period of 250 ms before response to 250 ms after response). A linear mixed-effects model with an imposed AR1 correlation structure was applied with power in the amygdala or hippocampus as the dependent variable, power in the motor cortex as the independent variable, and age and sex as covariates. A significant positive correlation between beta power in the motor cortex and beta power in the (A) amygdala (*p* < 0.001) and (B) hippocampus (*p* < 0.001) was observed. Additionally, a significant positive correlation between gamma power in the motor cortex and gamma motor in the (C) amygdala (*p* < 0.001) and (D) hippocampus (*p* < 0.001) was observed.

### Power differences between correct go and correct stop trials

In accordance with evidence of mesolimbic modulation of movement and prior imaging studies revealing mesolimbic contributions to motor inhibitory control, we hypothesized that the hippocampus and amygdala are similarly involved in response inhibition. Therefore, we first performed cluster-based permutation testing to identify specific time clusters in various frequency bands that were associated with stopping as compared to going in the motor cortex. We identified two significant clusters in the motor cortex for which power during correct go and stop responses differed. As seen in Figure 5A, cluster-based permutation testing identified significant differences from 390 ms to 740 ms after go signal presentation (*p* = 0.01) for delta power and from 360 ms to 730 ms after go signal presentation (*p* = 0.005) for theta power. During these two temporal windows, correct stop trials were associated with increased theta and delta power. We then evaluated whether delta and theta power in these time intervals were also significantly different between going and stopping in the amygdala and hippocampus. Comparing average delta power during correct go trials with average delta power during correct stop trials from 390 ms to 740 ms after go signal presentation, we found a significant difference for limbic regions (Wilcoxon-signed rank test, amygdala: *p* = 0.017, *z* = −2.85; hippocampus: *p* = 0.003, *z* = −3.37), with stopping also associated with increased average delta power in limbic regions. Similarly, comparing average theta power during these two trial types from 360 ms to 730 ms after go signal presentation revealed a significant difference for the amygdala (Wilcoxon-signed rank test: *p* = 0.018, *z* = −2.85), with correct stop trials similarly associated with higher average theta power. The difference in theta power in the hippocampus between going and stopping did not survive correction for multiple comparisons. Figure 5B displays these comparisons.

**Figure 5.**
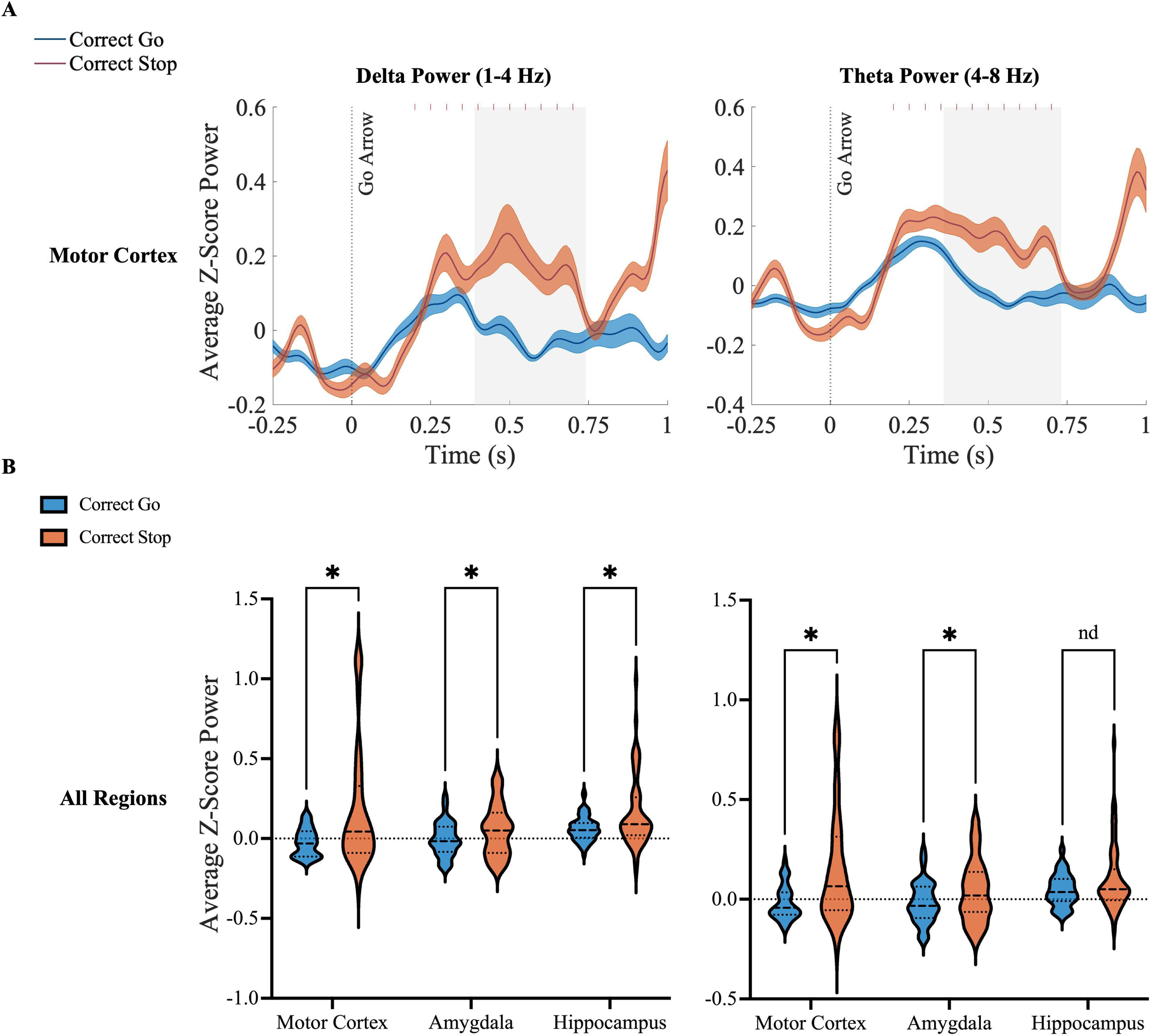
Power differences between correct go and correct stop trials. (A) Time courses of z-scored power during correct go and correct stop trials in the motor cortex for delta (1-4 Hz) and theta (4-8 Hz) power. Significant differences were found for delta (*p* = 0.01, 390 ms to 740 ms after go arrow onset) and theta power (*p* = 0.005, 360 ms to 730 ms after go arrow onset). Red ticks represent times during which stop signals could have occurred. Shading around lines indicates 99% confidence intervals and gray shaded boxes indicate time range of clusters that were significantly different between correct go and correct stop trials. (B) Violin plots of z-scored power in the delta and theta frequency ranges during the time periods that were significantly different between correctly making a response and correctly inhibiting a response. Delta power was significantly different in the amygdala (*p* = 0.017) and hippocampus (*p* = 0.003). Theta power was significantly different in the amygdala (*p* = 0.018).

### Power differences between correct and incorrect stop trials

We also evaluated neurophysiology correlates of response inhibition by comparing average power during correct and incorrect stop trials aligned to the stop arrow. Again, we determined the time windows significant for the motor cortex and limited our analyses for the amygdala and hippocampus to these periods. As shown in Figure 6A, we found significant differences in the motor cortex from 790 ms to 1330 ms after stop arrow appearance (*p* = 0.005) for delta power, from 760 ms to 1020 ms (*p* = 0.035) for theta power, from 1500 to 1720 ms (*p* = 0.005) for alpha power, from 40 ms to 240 ms (*p* = 0.02) for beta power, and from 270 ms to 1570 ms (*p* = 0.005) for gamma power. During these time intervals, incorrect stop trials were correlated with increased delta, theta, beta, and gamma power and decreased alpha power. Comparing average delta power during correct stop trials with average delta power during incorrect stop trials within the significant time window revealed a significant difference for the hippocampus (Wilcoxon-signed rank test: *p* = 5.8×10^−4^, *z* = 4.02), with failing to inhibit associated with decreased average delta power. Additionally, average alpha and beta power were significantly different for correct and incorrect stop trials in the amygdala (Wilcoxon-signed rank test, alpha power: *p* = 0.0013, *z* = 3.83, beta power: *p* = 2.1×10^−4^, *z* = 4.25), with incorrectly making a motor response correlated with reduced power. Finally, both the amygdala and hippocampus showed significant differences for average gamma power between the two trial types (Wilcoxon-signed rank test, amygdala: *p* = 3.3×10^−6^, *z* = −5.11, beta power: *p* = 1.8×10^−5^, *z* = −4.77), with incorrect stop trials associated with increased power as was seen in the motor cortex. No significant differences between correct and incorrect stop trials were found for theta power in the limbic areas. Figure 6B displays these comparisons.

**Figure 6.**
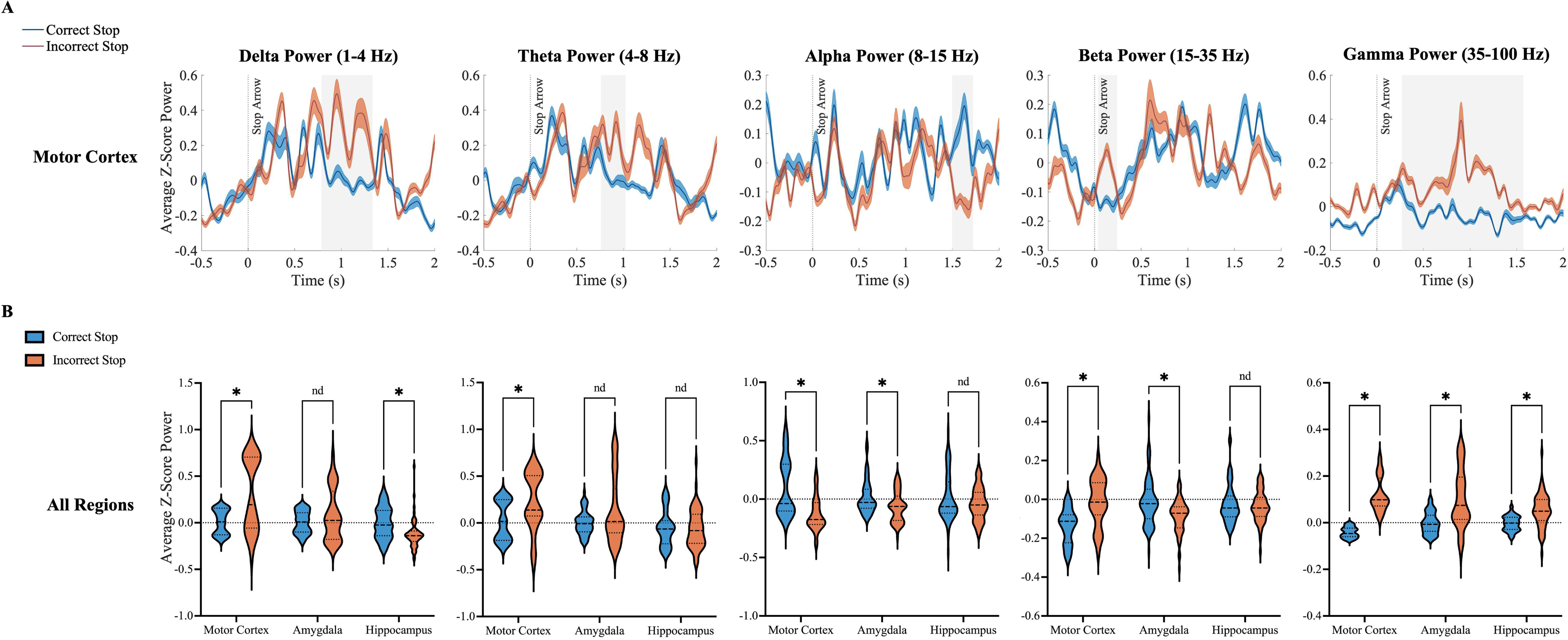
Power differences between correct and incorrect stop trials. (A) Time courses of z-scored power during correct and incorrect stop trials in the motor cortex separated by frequency band. Significant differences were found for delta power (*p* = 0.005, 790 ms to 1330 ms after stop arrow onset), theta power (*p* = 0.035, 760 ms to 1020 ms after stop arrow onset), alpha power (*p* = 0.005, 1500 to 1720 ms after stop arrow onset), beta power (*p* = 0.02, 40 ms to 240 ms after stop arrow onset), and gamma power (*p* = 0.005, 270 ms to 1570 ms after stop arrow onset). Shading around lines indicates 99% confidence intervals and gray shaded boxes indicate time range of clusters that were significantly different between correct and incorrect stop trials. (B) Violin plots of z-scored LFP power in all frequency bands during each respective significant time period while correctly inhibiting a response and incorrectly executing a response. Delta power was significantly different in the hippocampus (*p* < 0.001). Alpha power was significantly different in the amygdala (*p* = 0.0013). Beta power was significantly different in the amygdala (*p* < 0.001). Gamma power was significantly different in the amygdala (*p* < 0.001) and hippocampus (*p* < 0.001). Theta power was not significantly different in the limbic areas.

## Discussion

This study investigated the relationship between limbic regions and motor output by examining neural oscillatory dynamics during the stop-signal task. An analysis of beta and gamma power during motor response and baseline fixation revealed significant decreases in beta and gamma power in mesial temporal areas during movement planning and execution. Additionally, we found evidence of a correlation between limbic beta and gamma power and motor cortex beta and gamma power during movement execution. An exploratory analysis demonstrated significant differences between going and stopping for delta power in the amygdala and hippocampus and theta power in the amygdala. Finally, we found significantly elevated gamma oscillatory power in mesolimbic regions after participants failed to inhibit their responses, potentially representing an error monitoring signal.

### Relationships between beta and gamma power in mesolimbic regions and movement

The finding of reduced beta power in the motor cortex during a go response is in agreement with the well-established role of beta activity in movement. Beta oscillations in motor brain regions are suppressed during motor planning and execution. Conversely, beta activity in motor circuitry is elevated in Parkinson’s disease and is hypothesized to be anti-kinetic in nature.^13,29–32^ However, associations between movement and beta oscillatory activity in limbic regions have not been reliably shown. Recent sEEG studies using a direct-reach paradigm and a Go/No-Go task describe decreases in hippocampal beta power during the response period.^33,34^ Our results of reductions in limbic beta oscillatory power with motor response substantiate these conclusions for the hippocampus and expand upon them by establishing novel corresponding signal changes for the amygdala. The finding of beta suppression in the amygdala and hippocampus during motor planning and completion time periods supports a function for these nontraditional motor regions in movement. The reduction in beta activity appears to develop first and to a much larger degree in the motor cortex than in the mesolimbic areas, suggesting that the amygdala and hippocampus may serve as secondary modulators of action.

While we did not observe significant movement-related gamma power change in the motor cortex, previous work has demonstrated increased gamma band activity during movement.^18–20,30^ The opposing associations between beta and gamma power and movement allude to a movement-suppressing beta and movement-enhancing gamma model.^35,36^ Interestingly, we found that in the amygdala and hippocampus, gamma power decreased instead of increasing with motor response generation. A recent study of extracellular recordings in the rodent basolateral amygdala noted a spike in gamma activity immediately prior to active behavior initiation.^37^ Moreover, the study captured a considerable decrease in gamma oscillations following a conditioned response behavior and postulated that the reduction could be due to weakened synaptic firing.^37^ Our results seem to capture an overall reduction in gamma power from baseline during motor planning and execution, suggesting involvement of the limbic regions in movement. Prior work has suggested that in addition to contributing to action itself, hippocampal gamma power is likely also essential to the representation of movement trajectories.^38^

Furthermore, our findings of a positive relationship between beta and gamma power in the amygdala and hippocampus and beta and gamma power in the motor cortex indicate that limbic oscillations align with motor oscillations during movement. These correlations reinforce the idea that mesolimbic regions may behave in tandem with classic motor areas to generate action.

### Oscillatory power associated with motor inhibition

We found delta power increases in motor cortex when response was correctly inhibited during stop trials, with this same increase also seen in the amygdala and hippocampus. These results are in line with previous work using scalp EEG which found delta and theta band activity during motor inhibition.^39–41^ Although previously these oscillatory inhibitory signals were solely identified in midfrontal regions, our findings reveal that they similarly arise in mesial temporal nodes. In the amygdala and motor cortex, we also observed increased theta power during stopping. It is likely that theta and delta oscillations rise synchronously during motor adjustment.^40^ A non-human primate study of dorsal premotor area LFPs during a memory motor task with interruptions indicated that theta power is important for motor plan retrieval, perhaps suggestive of its function in inhibitory control.^42^ Our finding of delta and theta activity in limbic areas during response inhibition and the understanding that these slow waves are regularly associated with response inhibitory processes supports mesolimbic modulation of action control.

### Alpha and gamma power linked to error signals

We detected decreases in alpha power after incorrect stop trials in the amygdala and motor cortex, suggesting that the amygdala may influence post-error functioning. Prior investigations of error-related oscillations have similarly reported cortical alpha power reductions following errors relative to correct responses.^43–46^ This alpha suppression has been theorized to reflect augmented cortical arousal following a mistake due to the error commission itself or the infrequency of incorrect stop trials.^45,46^ It is possible that the traditionally “emotional” limbic system may impact behavior following undesirable performance.

Moreover, the discovery of prominent gamma oscillatory power increases in the amygdala, hippocampus, and motor cortex after unsuccessful stopping may further signify a role for these regions in error processing. Importantly, such increases were not seen in correct go trials similarly characterized by motor response, suggesting that this signal is specific to the error condition. Previous reports of LFP recordings during error monitoring described comparable increased broadband gamma power in the anterior insula and hippocampus.^47,48^ Our study reproduces this error-related gamma response in the hippocampus and reveals the same outcome for the amygdala and motor cortex. It is conceivable that high gamma power activity, specifically in the hippocampus, following error commission is intertwined with gamma’s assumed role in working memory.^48–50^

Our results demonstrate a relationship between motor output and neural oscillatory activity in the amygdala and hippocampus, supporting the concept of mesolimbic regulation of movement. There are several limitations of our study. We acknowledge that our sample size of 14 patients is relatively small and are optimistic that future studies with a larger number of participants will further validate our results. Moreover, it is possible that epilepsy patients may have altered brain activity. However, we intentionally removed channels from regions of seizure onset for our analyses in order to minimize potential confounds. Additionally, we recognize that the movement-related oscillatory findings indicate a link between limbic regions and movement but do not establish causality.

