## Supplementary Table 1 for "Mesolimbic local field potentials are modulated by motor control"

Stop-signal task performance

| **Participant ID** | **SSRT (ms)** | **GoRT (ms)** |
| --- | --- | --- |
| 1 | 295.86 | 766.46 |
| 2 | 298.25 | 815.84 |
| 3 | 198.88 | 684.36 |
| 4 | 285.60 | 452.75 |
| 5 | 276.54 | 483.41 |
| 6 | 178.21 | 595.90 |
| 7 | 331.68 | 625.43 |
| 8 | 212.92 | 497.63 |
| 9 | 230.53 | 561.46 |
| 10 | 295.76 | 678.72 |
| 11 | 382.27 | 758.89 |
| 12 | 358.33 | 804.02 |
| 13 | 311.46 | 589.49 |
| 14 | 345.40 | 448.42 |
| Average | 285.84 | 625.91 |
